# CD8 coreceptor-mediated focusing can reorder the agonist hierarchy of peptide ligands recognized via the T cell receptor

**DOI:** 10.1101/2020.09.23.310375

**Authors:** Mathew Clement, Lea Knezevic, Tamsin Dockree, James E. McLaren, Kristin Ladell, Kelly L. Miners, Sian Llewellyn-Lacey, Ore Francis, Andrew K. Sewell, John S. Bridgeman, David A. Price, Hugo A. van den Berg, Linda Wooldridge

## Abstract

CD8^+^ T cells are inherently cross-reactive and recognize numerous peptide antigens in the context of a given major histocompatibility complex class I (MHCI) molecule via the clonotypically expressed T cell receptor (TCR). The lineally expressed coreceptor CD8 interacts coordinately with MHCI at a distinct and largely invariant site to slow the TCR/peptide-MHCI (pMHCI) dissociation rate and enhance antigen sensitivity. However, this biological effect is not necessarily uniform, and theoretical models suggest that antigen sensitivity can be modulated in a differential manner by CD8. We used an intrinsically controlled system to determine how the relationship between the TCR/pMHCI interaction and the pMHCI/CD8 interaction affects the functional sensitivity of antigen recognition. Our data show that modulation of the pMHCI/CD8 interaction can reorder the agonist hierarchy of peptide ligands across a spectrum of affinities for the TCR.

**SIGNIFICANCE:** Sufficient immune coverage of the peptide universe within a finite host requires highly degenerate T cell receptors (TCRs). However, this inherent need for antigen cross-recognition is associated with a high risk of autoimmunity, which can only be mitigated by a process of adaptable specificity. We describe a mechanism that resolves this conundrum by allowing individual clonotypes to focus on specific peptide ligands without alterations to the structure of the TCR.

## INTRODUCTION

CD8^+^ T cells are critical for protective immunity against intracellular pathogens and various tumors. At the molecular level, activation is triggered by foreign or mutated peptide fragments presented on the cell surface by major histocompatibility complex class I (MHCI) molecules, which act as ligands for the somatically rearranged T cell receptor (TCR) and the germline-encoded coreceptor CD8 [1, 2]. The clonotypically expressed TCR confers antigen specificity by interacting with the peptide-binding platform of MHCI, which comprises the α1 and α2 domains, whereas the lineally expressed coreceptor CD8 is known to enhance antigen sensitivity by interacting primarily with the α3 domain of MHCI [3-7]. This latter interaction is biophysically and spatially independent of peptide-MHCI (pMHCI) engagement via the TCR [8]. However, the largely invariant nature of the pMHCI/CD8 interaction does not necessarily translate into a uniform gain of function, and theoretical studies have suggested that antigen sensitivity can be modulated in a differential manner, potentially altering the agonist hierarchy of peptide ligands for any given TCR [9, 10].

The pMHCI/CD8 interaction slows the dissociation rate of the TCR/pMHCI interaction [9, 11]. Functional sensitivity depends non-monotonically on this dissociation rate [12], as long as the system is limited by MHCI [10, 13, 14]. The nature of this relationship implies that functional sensitivity reaches a maximum at a particular dissociation rate. Strong agonists are relatively insensitive to modulation of the dissociation rate, because the curve has a negligible slope in the vicinity of the optimal value. In contrast, weak agonists are typically characterized by faster dissociation rates, modulation of which markedly alters functional sensitivity [15]. Accordingly, the pMHCI/CD8 interaction generally acts to increase agonist potency, maximizing the number of peptide ligands that can be recognized via a given TCR. However, theoretical models predict that ligands with dissociation rates below or close to the optimal value will respond differently, amounting to a differential focusing effect, whereby strong agonists can become less potent at dissociation rates beyond the optimal value. If operative *in vivo*, such an effect could allow individual clonotypes to focus on salient ligands [9], reconciling the inherent need for cross-reactivity with the inherent need for specificity [16].

We used a monoclonal system incorporating biophysically defined peptide ligands and variants of MHCI with altered coreceptor-binding properties to test the differential focusing hypothesis experimentally. In line with earlier predictions, we found that modulation of the pMHCI/CD8 interaction reordered the agonist hierarchy of peptide ligands recognized via the TCR.

## MATERIALS AND METHODS

### Cells

MEL5 TCR^+^ CD8^+^ J.RT3-T3.5 cells were maintained in RPMI 1640 medium containing 100 U/ml penicillin, 100 mg/ml streptomycin, 2 mM L-glutamine, and 10% heat-inactivated fetal calf serum (all from Thermo Fisher Scientific) (R10). Clonal MEL5 CD8^+^ T cells were maintained in R10 supplemented with 200 IU/ml IL-2 and 25 ng/ml IL-15 (both from PeproTech). The MEL5 TCR is specific for the heteroclitic HLA-A*0201-restricted Melan-A epitope ELAGIGILTV_26–35/A27L_ [17, 18]. HEK 293 cells were grown in Dulbecco’s Modified Eagle Medium (Sigma-Aldrich) supplemented with 100 U/ml penicillin, 100 mg/ml streptomycin, 2 mM L-glutamine, 10% heat-inactivated fetal calf serum, and 10 mM HEPES (all from Thermo Fisher Scientific). Hmy C1R cells expressing HLA-A*0201 (abbreviated from hereon as HLA-A2) and variants thereof were generated and maintained as described previously [19].

### Peptides

All peptides were synthesized at >95% purity using standard Fmoc chemistry (BioSynthesis Inc.).

### Lentiviruses

The α and β chains of the MEL5 TCR were engineered to contain mouse constant domains [20] and cloned into a single pSF-Lenti-EF1α lentiviral vector (Oxford Genetics) separated by an internal ribosomal entry site (IRES) sequence (Genewiz). The α and β chains of CD8 were cloned similarly into a single pSF-Lenti-EFα lentiviral vector (Oxford Genetics) separated by an IRES sequence (Genewiz). HEK 293 cells were cotransfected with the MEL5 TCR or CD8αβ lentiviral vectors and the packaging plasmids pMDLg/pRRE, pRSV-Rev, and pCMV-VSV-G using Turbofect Transfection Reagent (Thermo Fisher Scientific). Lentiviral particles were concentrated using Lenti-X Concentrator (Takara Bio).

### Generation of MEL5 TCR^+^ CD8^+^ J.RT3-T3.5 cells

TCR-deficient J.RT3-T3.5 cells were transduced with MEL5 TCR lentiviral particles and magnetically enriched using anti-murine TCRβ–PE (clone REA318) in conjunction with anti-PE MicroBeads (Miltenyi Biotec). MEL5 TCR^+^ J.RT3-T3.5 cells were then transduced with CD8αβ lentiviral particles, and MEL5 TCR^+^ CD8^+^ J.RT3-T3.5 cells were flow-purified using an Influx Cell Sorter (BD Biosciences).

### Quantification of activation-induced CD69

C1R cells expressing HLA-A2 D227K/T228A, wildtype HLA-A2, or HLA-A2 A245V/K^b^ were pulsed for 1 h with various concentrations of the peptides ELTGIGILTV (3T), ELAGIGILTV (ELA), or FATGIGIITV (FAT). Cells were then washed twice with RPMI 1640 medium containing 100 U/ml penicillin and 100 mg/ml streptomycin and resuspended in R10. Each assay included 1.5 × 10^5^ peptide-pulsed C1R cells and 5 × 10^4^ MEL5 TCR^+^ CD8^+^ J.RT3-T3.5 cells. Unpulsed targets were used as negative controls. Expression of CD69 on the surface of MEL5 TCR^+^ CD8^+^ J.RT3-T3.5 cells was measured after 6 h using the following directly conjugated monoclonal antibodies: anti-CD8α–PE-Cy7 (clone RPA-T8; Thermo Fisher Scientific), anti-CD8β–eFluor660 (clone SIDI8BEE; Thermo Fisher Scientific), anti-CD69–BV421 (clone FN50; BioLegend), and anti-HLA-A2–FITC (clone BB7.2; BioLegend). Non-viable cells were excluded from the analysis using LIVE/DEAD Fixable Aqua (Thermo Fisher Scientific). Data were acquired using a NovoCyte Flow Cytometer (ACEA Biosciences) and analyzed using FlowJo software version 10.6.1 (FlowJo LLC).

### Quantification of activation-induced IFN-γ

C1R cells expressing HLA-A2 D227K/T228A, wildtype HLA-A2, or HLA-A2/K^b^ were pulsed for 1 h with various concentrations of the peptides 3T, ELA, or FAT. Cells were then washed twice with RPMI 1640 medium containing 100 U/ml penicillin and 100 mg/ml streptomycin and resuspended in R10. Each assay included 6 × 10^4^ peptide-pulsed C1R cells and 3 × 10^4^ clonal MEL5 CD8^+^ T cells. Unpulsed targets were used as negative controls. Supernatants were harvested after 4 h and assayed for IFN-γ via ELISA (R&D Systems).

### Statistics

Functional assay data were processed using simultaneous non-linear least-squares parameter estimation encoded in *Mathematica* [21]. Functional sensitivity (*p*EC_50_) was expressed as the decimal cologarithm *p* of the 50% efficacy concentration (EC_50_). Data were analyzed using a one-way ANOVA or a two-way ANOVA with Tukey’s post-hoc test in *Mathematica* or Prism software version 8 (GraphPad).

## RESULTS

We used an intrinsically controlled system to determine how CD8 affects functional responses initiated via the MEL5 TCR. Ligand recognition in this system has been characterized previously using surface plasmon resonance [18, 22]. Biophysically defined peptide ligands, including a weak agonist (3T), the wildtype peptide (ELA), and a superagonist (FAT), were selected for the purposes of this work to introduce a range of TCR/pMHCI affinities (Table 1). C1R cells expressing HLA-A2 D227K/T228A, which abrogates the coreceptor interaction [23], wildtype HLA-A2, HLA-A2 A245V/K^b^, which enhances the coreceptor interaction [24], or HLA-A2 K^b^, which superenhances the coreceptor interaction [25], were used in parallel to introduce a range of pMHCI/CD8 affinities (Table 2). Importantly, surface plasmon resonance experiments have shown that none of these mutations, namely D227K/T228A, A245V/K^b^, and K^b^, affect the TCR/pMHCI interaction [11, 24].

**Table 1.**
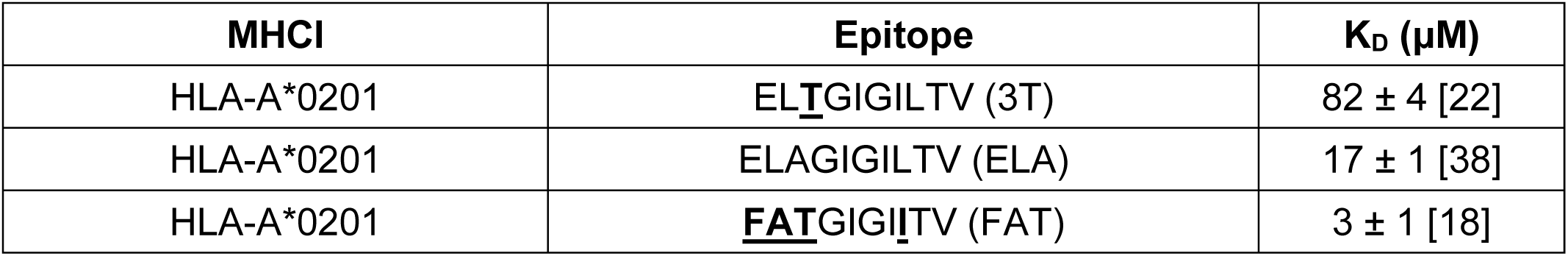
TCR/pMHCI dissociation constants for agonists of the MEL5 TCR.

**Table 2.** pMHCI/CD8 dissociation constants for variants of HLA-A*0201. NA: not applicable.

| **MHCI** | **Mutation** | **K<sub>D</sub> (μM)** |
| --- | --- | --- |
| HLA-A*0201 D227K/T228A | MHCI α3 domain | >10,000 [11] |
| HLA-A*0201 WT | NA | 137 ± 9.7 [11] |
| HLA-A*0201 A245V/K<sup>b</sup> | MHCI α3 domain | 27 ± 1 [11] |
| HLA-A*0201 K<sup>b</sup> | MHCI α3 domain | 11 [24] |

In preliminary experiments, we quantified CD69 on the surface of MEL5 TCR^+^ CD8^+^ J.RT3-T3.5 cells as a measure of activation in response to 3T, ELA, or FAT presented in the context of HLA-A2 D227K/T228A, wildtype HLA-A2, or HLA-A2 A245V/K^b^ (Supplementary Figures S1 & S2). Functional sensitivity was determined as the *p*EC_50_ value for each parameter combination (Figure 1A). In the absence of a pMHCI/CD8 interaction (HLA-A2 D227K/T228A), activation was a simple function of TCR/pMHCI affinity (Figure 1A and Supplementary Figures S1 & S2). The agonist potencies of 3T and ELA were enhanced in the context of HLA-A2 and HLA-A2 A245V/K^b^ relative to HLA-A2 D227K/T228A (Figure 1A). In contrast, the agonist potency of FAT was only marginally enhanced in the context of HLA-A2 relative to HLA-A2 D227K/T228A and, consistent with the notion of an optimal activation window, decreased slightly in the context of HLA-A2 A245V/K^b^ relative to HLA-A2 (Figure 1A & Supplementary Figure S3A). As a consequence, the agonist potency of FAT relative to the agonist potency of ELA was reduced at higher pMHCI/CD8 affinities (Figure 1B), and in three of four replicate experiments, ELA was the most potent ligand in the context of HLA-A2 and HLA-A2 A245V/K^b^ (Figure 1B–E).

**Figure 1.**
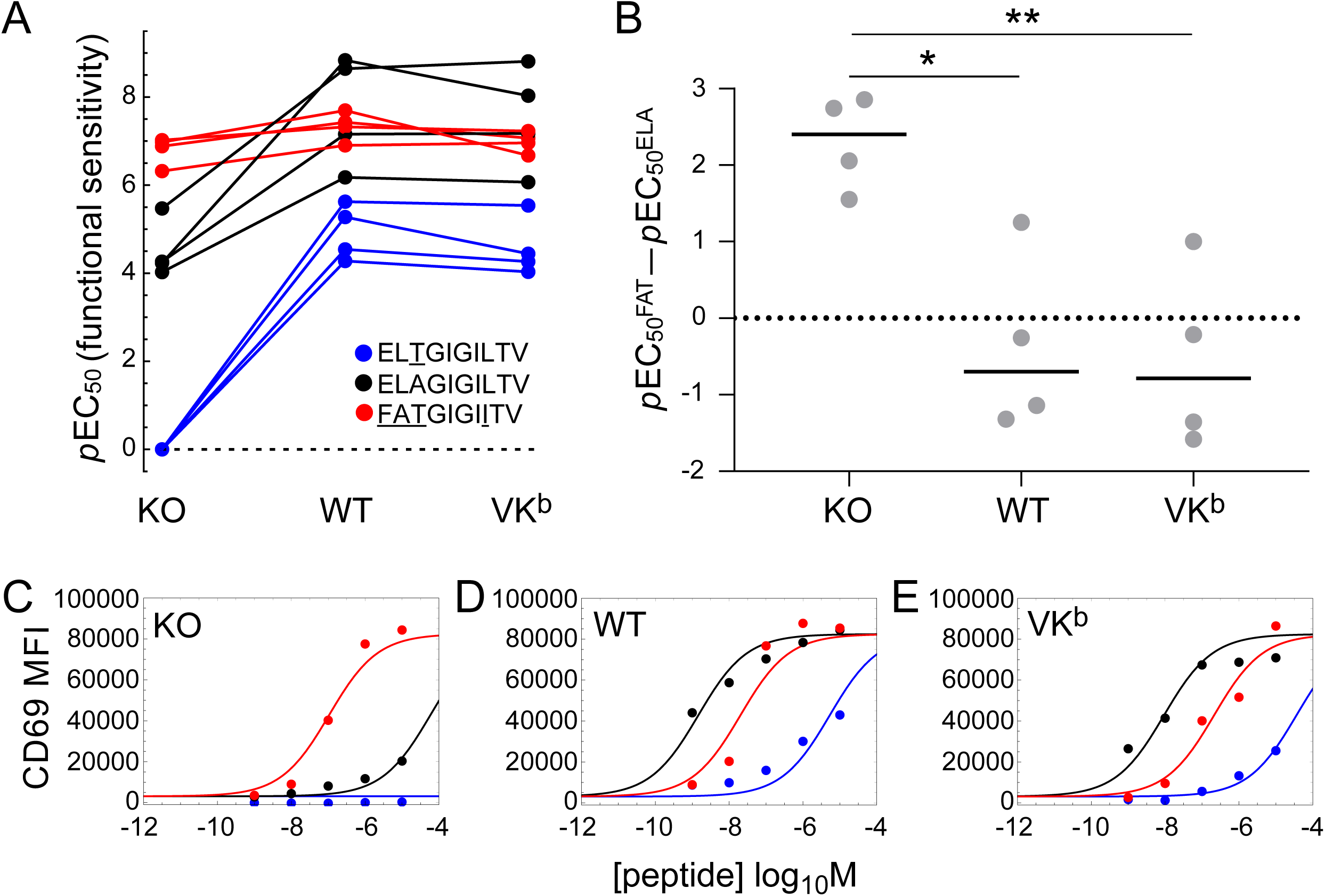
CD8 reorders the agonist hierarchy of peptide ligands that induce the expression of CD69. MEL5 TCR^+^ CD8^+^ J.RT3-T3.5 cells were activated for 6 h with C1R cells expressing HLA-A2 D227K/T228A (KO), wildtype HLA-A2 (WT), or HLA-A2 A245V/K^b^ (VK^b^) pulsed with various concentrations of 3T (blue), ELA (black), or FAT (red). Surface expression of CD69 was measured via flow cytometry. (**A**) Functional sensitivity (*p*EC50) for each peptide ligand in the context of each MHCI. Four replicate experiments are shown. The value for 3T in the context of HLA-A2 D227K/T228A was set to zero for graphical purposes and treated as missing data for statistical purposes. *P* < 0.0001 for the ligand effect, *P* < 0.0001 for the MHCI effect (two-way ANOVA with Tukey’s post-hoc test). (**B**) The agonist potency of FAT relative to the agonist potency of ELA expressed as *p*EC_50_^FAT^ − *p*EC_50_^ELA^, which is equivalent to the logarithm of the fold difference in functional sensitivity. Four replicate experiments are shown. Horizontal bars indicate median values. \**P* < 0.05, \*\**P* < 0.01 (one-way ANOVA with Tukey’s post-hoc test). (**C**–**E**) Representative peptide titration experiment used to calculate the parameters in A and B. Curves were fitted in *Mathematica*. All four replicate experiments are shown in Supplementary Figures S1 & S2.

To confirm these findings, we quantified the production of IFN-γ by clonal MEL5 CD8^+^ T cells in response to 3T, ELA, or FAT presented in the context of HLA-A2 D227K/T228A, wildtype HLA-A2, or HLA-A2 K^b^ (Supplementary Figures S4 & S5). Functional sensitivity was again determined as the *p*EC_50_ value for each parameter combination (Figure 2A). The activation data were largely analogous to those obtained with MEL5 TCR^+^ CD8^+^ J.RT3-T3.5 cells. In particular, the agonist potency of FAT was enhanced in the context of HLA-A2 relative to HLA-A2 D227K/T228A and decreased slightly in the context of HLA-A2 K^b^ relative to HLA-A2 (Figure 2A & Supplementary Figure S3B), mirroring the downturn in functional sensitivity observed with MEL5 TCR^+^ CD8^+^ J.RT3-T3.5 cells in the context of HLA-A2 A245V/K^b^ relative to HLA-A2 (Figure 1A & Supplementary Figure S3A). As a consequence, the agonist potency of FAT relative to the agonist potency of ELA was again reduced at higher pMHCI/CD8 affinities (Figure 2B), and in three of four replicate experiments, ELA was the most potent ligand in the context of HLA-A2 K^b^ (Figure 2B–E).

**Figure 2.**
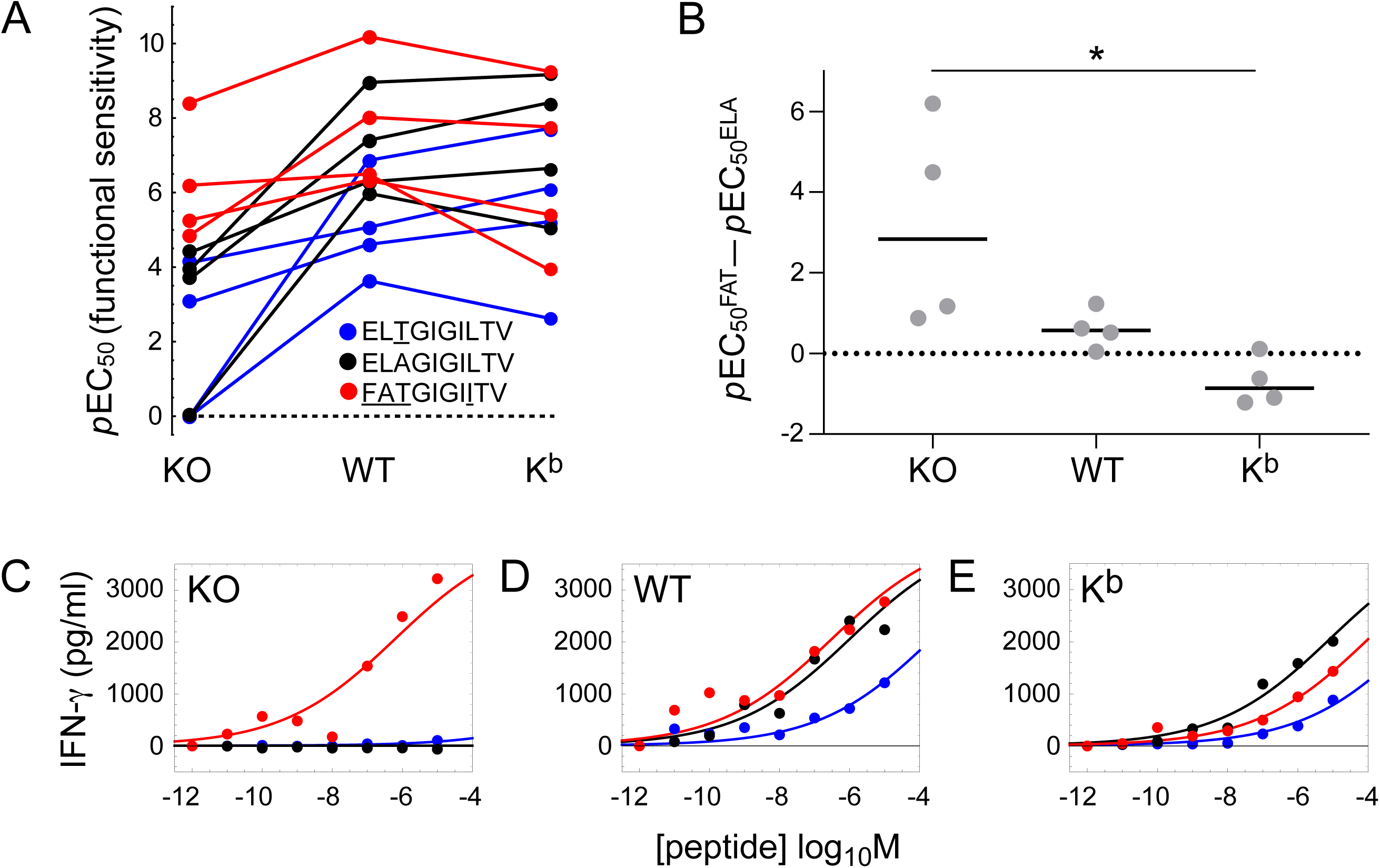
CD8 reorders the agonist hierarchy of peptide ligands that induce the production of IFN-γ. Clonal MEL5 CD8^+^ T cells were activated for 4 h with C1R cells expressing HLA-A2 D227K/T228A (KO), wildtype HLA-A2 (WT), or HLA-A2 K^b^ (K^b^) pulsed with various concentrations of 3T (blue), ELA (black), or FAT (red). Secretion of IFN-γ was measured via ELISA. (**A**) Functional sensitivity (*p*EC50) for each peptide ligand in the context of each MHCI. Four replicate experiments are shown. Values below the limit of estimation were set to zero for graphical purposes and treated as missing data for statistical purposes. *P* = 0.0042 for the ligand effect, *P* = 0.00069 for the MHCI effect (two-way ANOVA with Tukey’s post-hoc test). (**B**) The agonist potency of FAT relative to the agonist potency of ELA expressed as *p*EC_50_^FAT^ − *p*EC_50_^ELA^, which is equivalent to the logarithm of the fold difference in functional sensitivity. Four replicate experiments are shown. Horizontal bars indicate median values. \**P* < 0.05 (one-way ANOVA with Tukey’s post-hoc test). (**C**–**E**) Representative peptide titration experiment used to calculate the parameters in A and B. Curves were fitted in *Mathematica*. All four replicate experiments are shown in Supplementary Figures S4 & S5.

Collectively, these results can be interpreted and understood in biological terms if two key assumptions are made: (i) functional sensitivity depends non-monotonically on the TCR/pMHCI dissociation rate [12]; and (ii) the pMHCI/CD8 interaction affects the TCR/pMHCI dissociation rate by an invariant factor, equivalent to translation on a logarithmic scale (Figure 3). In this scenario, ligands that are recognized poorly in the absence of a pMHCI/CD8 interaction become more potent in the presence of a physiological pMHCI/CD8 interaction and achieve optimal agonist potency in the presence of a supraphysiological pMHCI/CD8 interaction, whereas ligands that are recognized strongly in the absence of a pMHCI/CD8 interaction straddle an optimum in the presence of a physiological pMHCI/CD8 interaction and become less potent in the presence of a supraphysiological pMHCI/CD8 interaction. Accordingly, the agonist hierarchy of peptide ligands, which is dictated in isolation by the TCR/pMHCI interaction, can be reordered as a function of coengagement by CD8.

**Figure 3.**
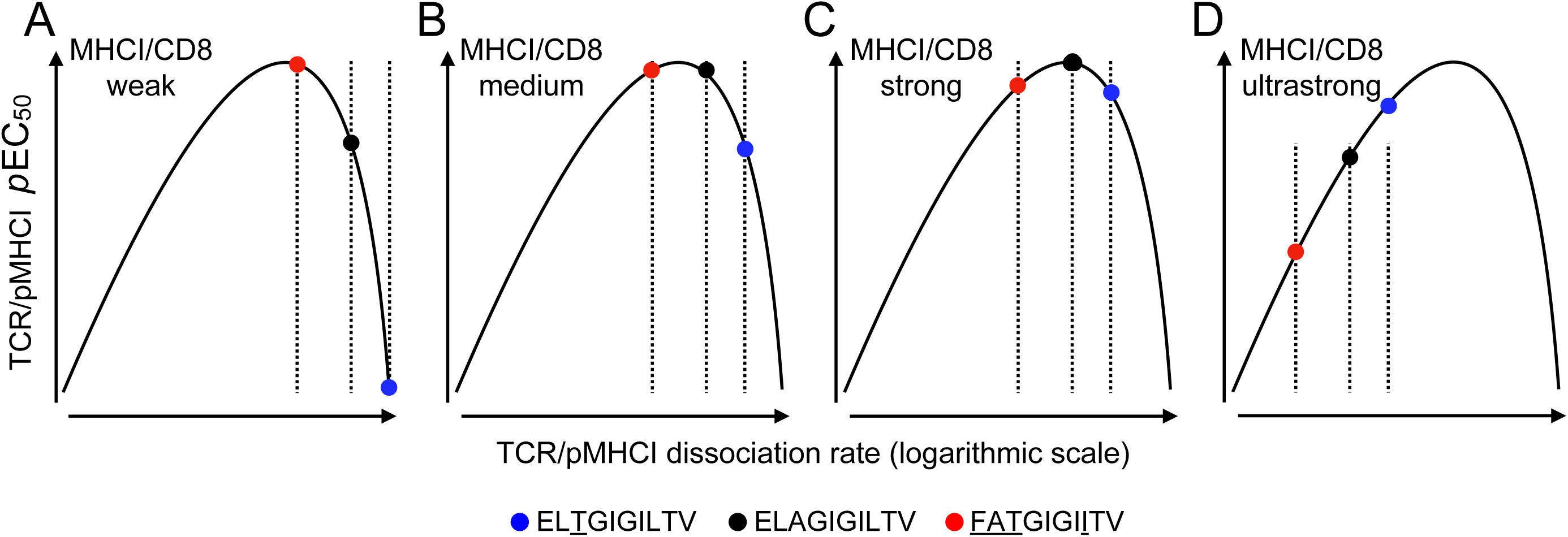
Theoretical interpretation of the differential focusing effect mediated by CD8. Graphical representation of the differential focusing effect based on two key assumptions: (i) functional sensitivity depends non-monotonically on the TCR/pMHCI dissociation rate; and (ii) the pMHCI/CD8 interaction affects the TCR/pMHCI dissociation rate by an invariant factor, equivalent to translation on a logarithmic scale. (**A**–**C**) Modulation of the pMHCI/CD8 interaction moves peptide ligands along this curve, altering the agonist hierarchy as a function of the TCR/pMHCI dissociation rate. (**D**) A hypothetical ultrastrong pMHCI/CD8 interaction would be expected to reverse the agonist hierarchy from FAT > ELA > 3T to FAT < ELA < 3T.

## DISCUSSION

CD8^+^ T cells are inherently promiscuous and can recognize more than a million different peptide ligands via the TCR [21, 26-28]. It is well established that CD8 can enhance the functional sensitivity of antigen recognition, but in any given monoclonal system, it does not necessarily follow that CD8 will affect the agonist potency of every cognate ligand in a similar manner. Indeed, theoretical studies have suggested that the agonist hierarchy of peptide ligands can be modified or even reversed across a range of pMHCI/CD8 affinities, such that a differential focusing effect acts to optimize the recognition of particular ligands in the context of an individual TCR [9, 10, 14]. Our data provide experimental confirmation of these predictions.

The biological relevance of differential focusing remains unknown, but hypothetical considerations suggest that such an effect may be advantageous *in vivo*, especially if accompanied by feedback mechanisms that enable the process of specificity adjustment to converge on a foreign antigen. Optimal recognition of a particular agonist in this manner would maximize immune efficacy during the process of clonal expansion and simultaneously minimize the risk of autoimmunity. Affinity maturation subserves an equivalent function in B cells. In more general terms, differential focusing also provides a solution to the “Mason paradox”, allowing a high degree of immune specificity alongside sufficient coverage of the peptide universe within a relatively small naive repertoire via the incorporation of degenerate TCRs [16].

Although it remains to be determined how differential focusing could operate *in vivo* and to what extent this might occur throughout the lifespan of any given clonotype, elegant studies have already provided important mechanistic clues. For example, double-positive thymocytes can transcriptionally downregulate CD8 [29], and antigen encounters in the periphery dynamically can modulate clonal responsiveness via the selective internalization of CD8 [30]. In addition, coreceptor use can be switched between the functionally distinct isoforms CD8αα and CD8αβ [31], which are further modifiable via glycosylation [32-34], and cytokine signals can transcriptionally alter the expression of CD8 [35]. All of these processes affect the signaling threshold for activation via the TCR in a manner akin to affinity variation in the pMHCI/CD8 interaction [11, 36]. Accordingly, functional sensitivity depends on the kinetics of signalosome development [9, 10], which is determined by agonist potency and regulated by CD8 [37].

In line with earlier theoretical predictions, the data presented here show that agonist potency, quantified in terms of functional sensitivity, can be differentially modulated across a range of TCR/pMHCI affinities by CD8. If this phenomenon occurs *in vivo*, as suggested by previous mechanistic studies, then immune reactivity could be focused on individual peptide ligands in the context of antigen-driven clonal expansions. On the basis of these collective observations, we propose that specificity adjustment operates at the level of individual clonotypes to safeguard the host in the face of an ongoing immune response, simultaneously facilitating the targeted delivery of effector functions and mitigating the risk of bystander damage, which can be triggered by inherently degenerate and therefore potentially autoreactive TCRs.

## ACKNOWLEDGEMENTS

We thank Andrew Herman and Lorena Sueiro Ballesteros for assistance with cell sorting at the University of Bristol Flow Cytometry Facility. This work was funded by the Wellcome Trust (WT079848MA and WT099067AIA). Additional support was received from the Horizon 2020 Research and Innovation Programme of the European Union under Marie Sklodowska-Curie Grant Agreement 721358 and the Biotechnology and Biological Sciences Research Council (Grant BB/H001085/1). A.K.S. and D.A.P. were supported by Wellcome Trust Senior Investigator Awards.

## AUTHOR CONTRIBUTIONS

M.C., L.K., T.D., J.E.M., K.L.M., S.L.L., and O.F. performed experiments. A.K.S. and J.S.B. provided key reagents. K.L., D.A.P., H.A.v.d.B., and L.W. supervised the research. D.A.P., H.A.v.d.B., and L.W. wrote the manuscript.

