## Supplementary Information for "CD8 coreceptor-mediated focusing can reorder the agonist hierarchy of peptide ligands recognized via the T cell receptor"

#### SUPPLEMENTARY FIGURE LEGENDS

**Supplementary Figure S1. CD8 reorders the agonist hierarchy of peptide ligands that induce the expression of CD69.** MEL5 $TCR^+$ $CD8^+$ J.RT3-T3.5 cells were activated for 6 h with C1R cells expressing HLA-A2 D227K/T228A (KO), wildtype HLA-A2 (WT), or HLA-A2 A245V/$K^b$ ($VK^b$) pulsed with various concentrations of 3T (blue), ELA (black), or FAT (red). Surface expression of CD69 was measured via flow cytometry. (**A–C**) Experimental replicate 1. (**D–F**) Experimental replicate 2. (**G–I**) Experimental replicate 3. (**J–L**) Experimental replicate 4.

**Supplementary Figure S2. CD8 reorders the agonist hierarchy of peptide ligands that induce the expression of CD69.** Experimental details as in Supplementary Figure S1. Curves were fitted to the same data in *Mathematica*.

**Supplementary Figure S3. Enhanced coreceptor interactions reduce the potency of a strong agonist recognized via the MEL5 TCR.** (**A**) Box and whisker plots summarizing the data shown in Figure 1A for FAT. (**B**) Box and whisker plots summarizing the data shown in Figure 2A for FAT.

**Supplementary Figure S4. CD8 reorders the agonist hierarchy of peptide ligands that induce the production of IFN-γ.** Clonal MEL5 $CD8^+$ T cells were activated for 4 h with C1R cells expressing HLA-A2 D227K/T228A (KO), wildtype HLA-A2 (WT), or HLA-A2 $K^b$ ($K^b$) pulsed with various concentrations of 3T (blue), ELA (black), or FAT (red). Secretion of IFN-γ was measured via ELISA. (**A–C**) Experimental replicate 1. (**D–F**) Experimental replicate 2. (**G–I**) Experimental replicate 3. (**J–L**) Experimental replicate 4. Each data point represents the mean of duplicate measurements. Error bars show SD.

**Supplementary Figure S5. CD8 reorders the agonist hierarchy of peptide ligands that induce the production of IFN-γ.** Experimental details as in Supplementary Figure S4. Curves were fitted to the same data in *Mathematica*.

### Figure S1

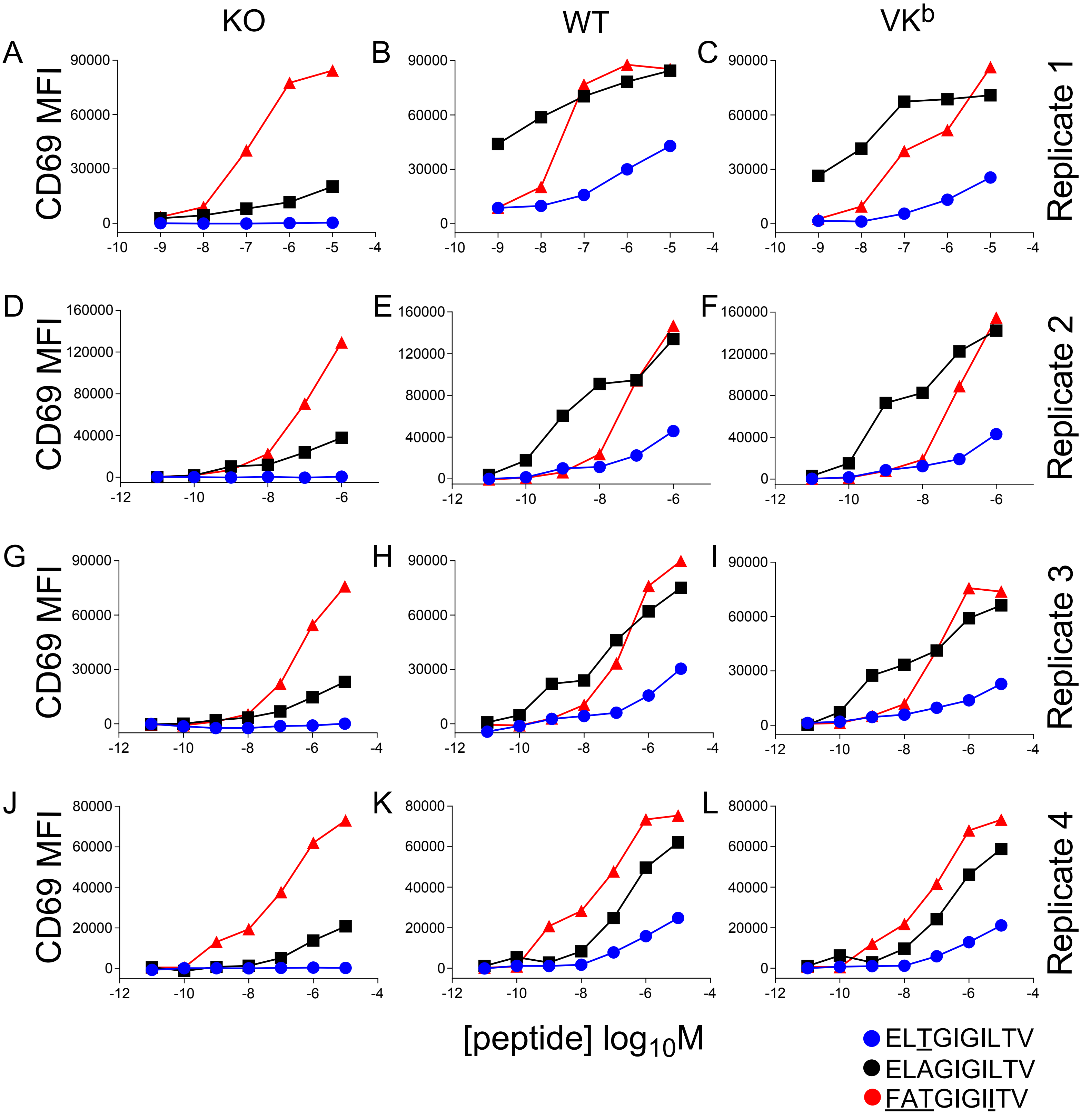

KO
WT
VK$^{b}$
A
B
C
D
E
F
G
H
I
J
K
L
CD69 MFI
Replicate 1
Replicate 2
Replicate 3
Replicate 4
[peptide] $\log_{10}$M
ELTGIGILTV
ELAGIGILTV
FATGIGIITV

### Figure S2

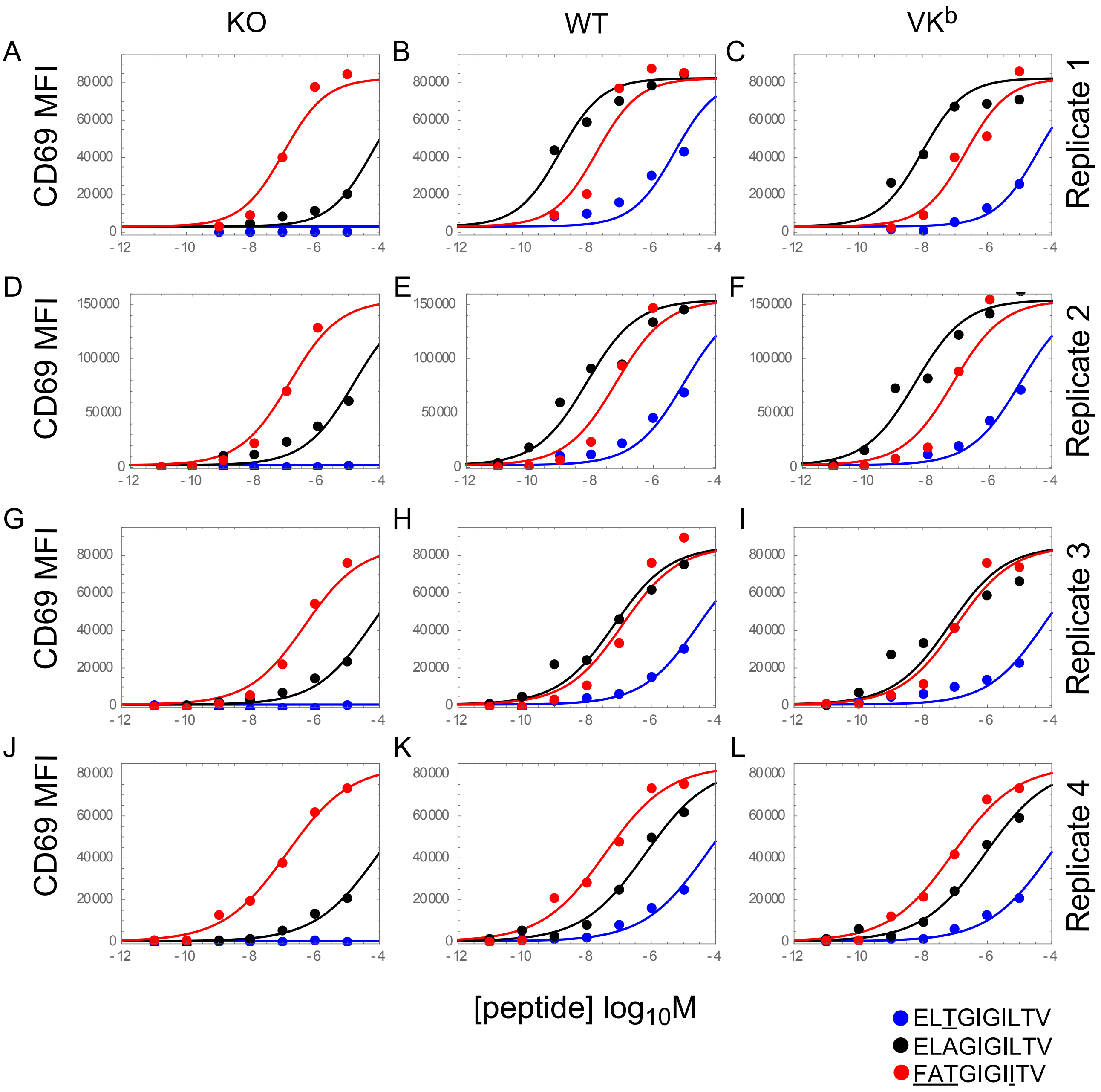

KO
WT
VK$^{b}$
A
B
C
D
E
F
G
H
I
J
K
L
CD69 MFI
Replicate 1
Replicate 2
Replicate 3
Replicate 4
0
20000
40000
60000
80000
50000
100000
150000
-12
-10
-8
-6
-4
[peptide] $log_{10}M$
ELTGIGILTV
ELAGIGILTV
FATGIGIITV

Figure S3

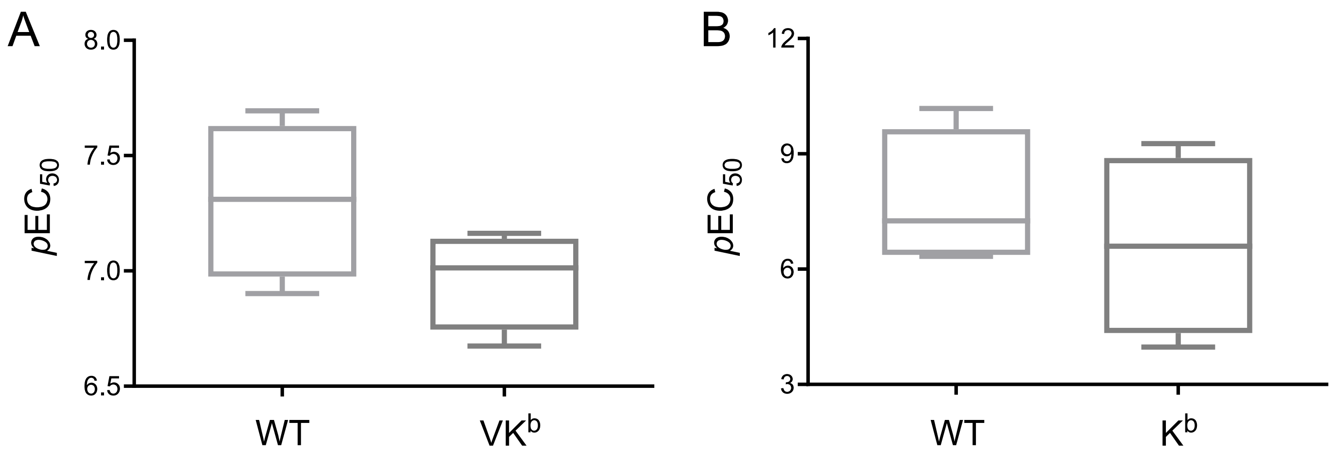

### Figure S4

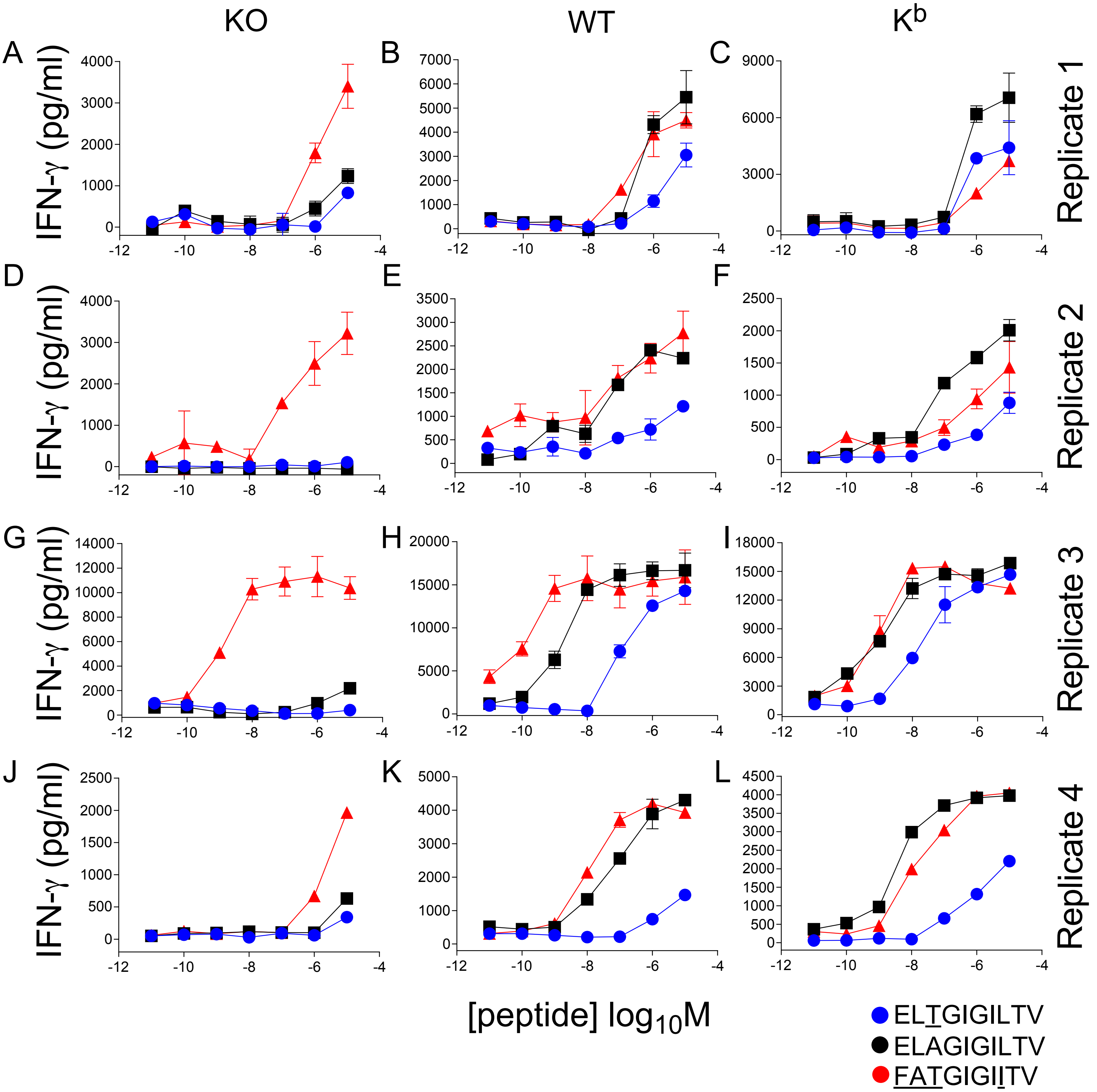
KO
WT
K^b
A
B
C
D
E
F
G
H
I
J
K
L
IFN-γ (pg/ml)
Replicate 1
Replicate 2
Replicate 3
Replicate 4
[peptide] log10M
ELTGIGILTV
ELAGIGILTV
FATGIGIITV

Figure S5

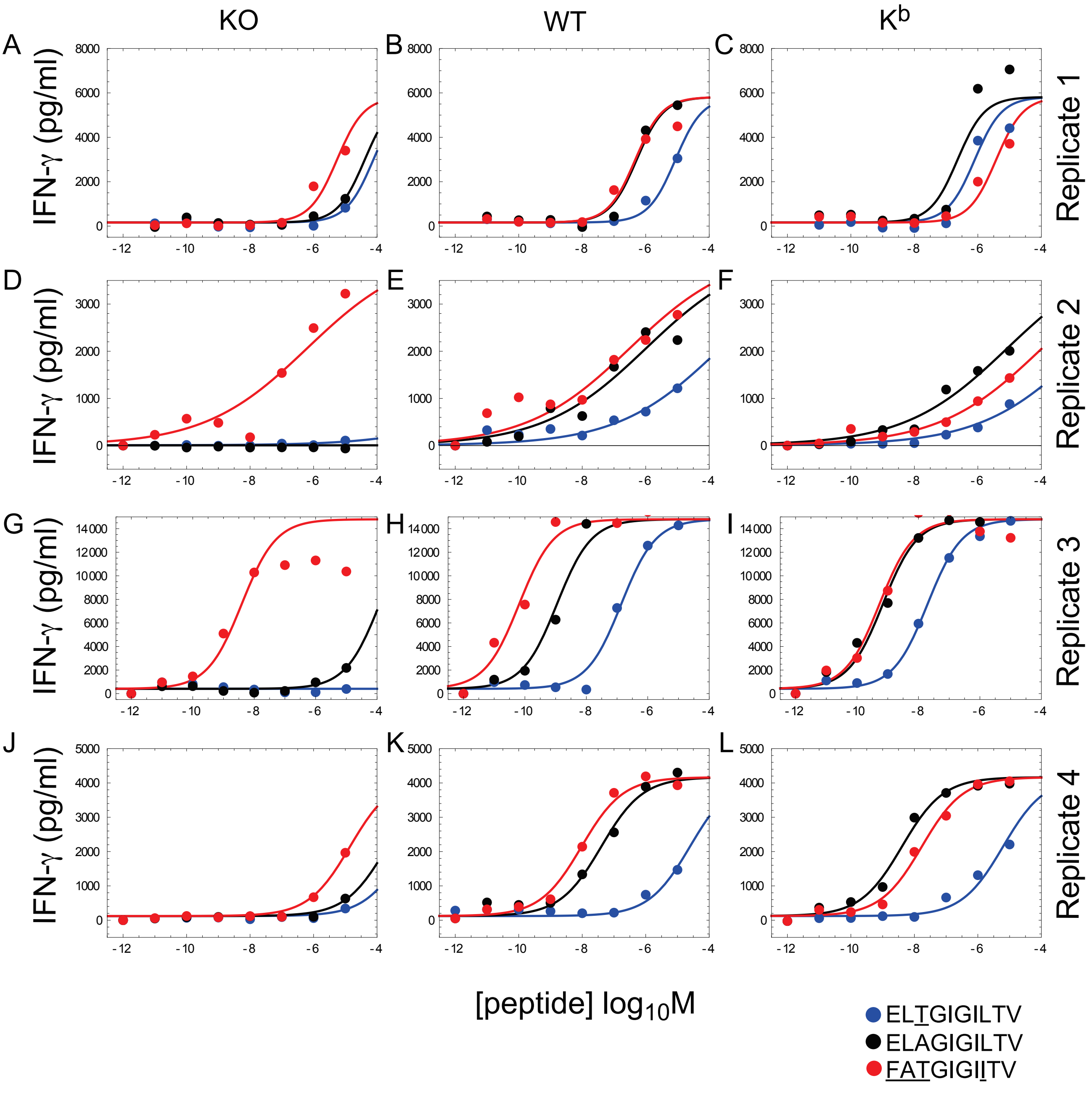

KO
WT
K^b
A
B
C
D
E
F
G
H
I
J
K
L
IFN-γ (pg/ml)
Replicate 1
Replicate 2
Replicate 3
Replicate 4
[peptide] log10M
ELTGIGILTV
ELAGIGILTV
FATGIGIITV
